# Renin is critical for Renin Lineage Cell Plasticity and Migration in experimental crescentic Glomerulonephritis

**DOI:** 10.64898/2026.02.11.705281

**Authors:** Shila Azizolli, Sagor Halder, Anne Steglich, Akua Annoh, Florian Gembardt, Irina Simonova, Jan Sradnick, Andreas Dahl, Rajinder Gupta, Stefan R. Bornstein, Vladimir Todorov, Hannah Weissbach, Christian Hugo

## Abstract

**Key Points:**

- Renin deficiency in renin-lineage cells worsened crescentic injury and impaired cell migration, revealing a protective role for renin in crescentic glomerulonephritis.
- Loss of renin shifted renin-lineage cells signaling toward interferon/STAT1-driven
- Renin-lineage cell ablation in crescentic glomerulonephritis induced a less inflammatory disease time-course.

**Background:** The adult juxtaglomerular renin-lineage cell (RLC) niche contributes to intraglomerular repair after injury, but their role in highly inflammatory crescentic glomerulonephritis (cGN) remains unclear. While angiotensin II–AT1R signaling promotes fibrosis and inflammation, the contribution of the RLCs, and of renin expression within RLCs, to cGN outcome has not been investigated.

**Methods:** We used tdTomato lineage-tracing to track RLCs in wild-type (WT) and renin-knockout (RenKO) mice following cGN induction. RLC migration and glomerular injury were quantified histologically. Single-cell RNA sequencing was performed on isolated tdTomato-positive cells at day 10 and 21 after injury to characterize transcriptional programs. Disease progression was additionally examined in mice with diphtheria toxin A–mediated (DTA) RLC ablation.

**Results:** RLCs were detected within injured glomeruli during cGN, with sporadic localization to crescentic lesions. Genetic renin deletion in RLCs worsened cGN outcomes, with RenKO mice developing increased albuminuria (by 306%), crescent formation (by 50%) and podocyte loss (by 15%) by day 21 compared to WT controls. Renin-deficient RLCs exhibited a reduced intraglomerular migratory response with decreased colocalization with mesangial and podocytes cell markers. Single-cell transcriptomic analysis supports an immunomodulatory reparative phenotype in WT RLCs. In contrast, RenKO RLCs displayed enrichment of interferon-stimulated genes and pathways suppressing cell migration. RLC ablation reduced macrophage infiltration, but did not alter disease progression, suggesting compensatory cellular mechanisms.

**Conclusions:** Renin expression supports the plasticity and injury-associated responses of RLCs during cGN. Loss of renin shifts RLCs toward an interferon-driven inflammatory and antimigratory phenotype that aggravates glomerular injury, while ablation of the RLCs may be compensated without major outcome changes.

## Introduction

Crescentic glomerulonephritis (cGN) represents a group of immune-mediated rapidly progressive kidney diseases arising from dysregulated immune responses^1^. Despite diverse etiologies, these conditions lead to disruption of the glomerular basement membrane (GBM), allowing macrophages, fibrin, and other immune and plasma constituents to breach Bowman’s space^2–4^. This triggers parietal epithelial cell (PEC) proliferation and crescent formation—the hallmark of cGN. Current therapies for cGN rely primarily on broad immunosuppression, which carries significant toxicity and often fails to prevent progression to end-stage kidney disease^5^. Understanding the cellular mechanisms driving crescent formation and identifying intrinsic renal cell populations that modulate local immune responses and repair could provide more specific therapeutic strategies with less side effects.

Renin-lineage cells (RLCs) represent a particularly relevant population in this context. During kidney development, RLCs originate from Foxd1⁺ stromal progenitor cells in the metanephric mesenchyme, differentiate into renin-expressing precursors that populate developing arteriolar walls throughout the kidney vasculature, and give rise to about 10% of all kidney cells ^6^. In adulthood, these cells become restricted to the juxtaglomerular apparatus (JGA), where they form a specialized precursor cell niche^7,8^.

Originally characterized for their role in the renin-angiotensin system (RAS), RLCs function as kidney-resident progenitors with demonstrated regenerative capacity. In several glomerular injury models, RLCs migrate from their juxtaglomerular niche and differentiate into multiple glomerular cell types. This transdifferentiation involves phenotypic switching characterized by loss of renin expression and acquisition of cell type-specific markers such as α8-integrin for mesangial cells (MC), WT1 or synaptopodin for podocytes, and claudin-1 for PECs^9–12^.

Beyond their established progenitor cell role, emerging evidence suggests RLCs may also participate in immune modulation. Although direct immune effector functions of juxtaglomerular RLCs remain to be demonstrated, renin-expressing B-1 lymphocytes exhibit immune regulatory functions distinct from their canonical roles^13,14^. Furthermore, the RAS—where renin is the rate-limiting enzyme—has well documented pro-inflammatory and pro-fibrotic effects in kidney disease^15,16^. Angiotensin II (AngII), the primary effector peptide of the RAS, promotes inflammatory cell recruitment, oxidative stress, and fibrosis through AT1 receptor signaling^17^. Conversely, alternative RAS-axes mediated by AT2 receptors and the ACE2-Ang (1-7)-Mas pathway exert protective, anti-inflammatory effects^18,19^. This duality raises the question of whether RLC-derived renin and local signaling influence immune-mediated injury in cGN, and whether RLC responses are determined by their renin production.

Despite their strategic positioning and capacity for regeneration and potential immune modulation, the role of RLCs in cGN remains unexplored. Whether RLCs respond to severe inflammatory injury by mobilizing for regeneration, participating in local immune responses, or both is unknown. Most critically, whether RLC-derived renin contributes to disease progression, a question with direct therapeutic implications, has not been addressed. To investigate RLC function and their broader role in cGN, we employed genetic lineage tracing to track their behavior and, either ablated adult juxtaglomerular RLCs or deleted renin from RLCs to determine whether RLC renin-production alters disease progression in cGN.

## Materials & Methods

### Animals

We used three inducible RPC-transgenic mouse strains. The control strain (mRen-rtTAm2-LC1-tdT, short WT) expresses the red fluorescent reporter tdTomato (tdT) in RPC ^20,21^. In the DTA strain (mRen-rtTAm2-LC1-tdT-DTA), induction triggers diphtheria toxin A expression in RPCs, leading to their ablation. In the RenKO strain (mRen-rtTAm2-LC1-tdT-Ren, RenKO), renin is selectively deleted in RPCs. The generation of the knockout mice is described in the supplement.

Animals were housed at 12:12-h light-dark cycle and had access to water and food ad libitum. All animal experiments were performed in accordance with the Federal Law on the Use of Experimental Animals in Germany and were approved by the local authorities. Animal experiments were consistent with the National Institutes of Health (NIH) Guide for the Care and Use of Laboratory Animals (NIH Pub. No. 85-23, Revised 2011).

### Experimental procedure

To induce tdTomato expression, male renin cell-ablated or renin knockout mice received doxycycline (2 mg/ml; scienTest-bioKEMIX GBR, Germany), enalapril (10 mg/kg; Ratiopharm, Germany), and sucrose (5%; Carl Roth GmbH & Co. KG, Germany) in drinking water for 16 days.

After one week washout, mice were injected intravenously via the tail vein with human TNFα (100 µg/kg body weight, 30 µg/ml in PBS). Two hours later, animals were injected intraperitoneally with anti-GBM serum (3 g/kg body weight, produced in sheep, 375 mg/ml in PBS). Control mice received PBS alone.

### Evaluation of kidney function

Glomerular filtration rate (GFR) was measured transdermally as previously described ^22,23^. Albumin (Biomol, #E99-134) and creatinine (Diazyme, #DZ072B-CAL) ELISA was performed according to the manufacturer’s instructions to calculate urinary albumin to creatine ratio.

Ten days after model induction unilateral nephrectomy was performed as previously described^21^. At the end of the experiment, mice were anesthetized by intraperitoneal injection of ketamine/xylazine (100 mg/kg bodyweight ketamine, 1 mg/kg bodyweight xylazine) and kidneys collected.

### Immunofluorescence staining

Kidney biopsies were fixed in zinc fixative (0.5 g calcium acetate, 5 g zinc acetate, 5 g ZnCl_2_ dissolved in 0.1 M TRIS-buffered solution (pH 7.4)) overnight at 4°C and processed as described^24^. Immunofluorescence staining was performed as previously described^6^ using antibodies listed in Supplemental Table 1. Sections were scanned using the AxioScanZ.1 (Zeiss, Germany) at the Light Microscopy Facility of the CMCB of Technical University Dresden.

Paraffin-embedded kidneys were cut into 2 µm thick sections, deparaffinized, rehydrated and stained with periodic acid–Schiff stain (PAS) and hematoxylin as counter stain. Automated evaluation of glomerular size and PAS-positive area was done as previously described^25^. Ten whole-kidney sections/group were manually counted for crescent formation, intraglomerular tdT-positive signal, number of glomerular WT1-positive and ERG-positive cells.

### RNA Isolation

Total RNA was extracted from kidney samples using TRIzol™ reagent (Invitrogen, USA) following the manufacturer’s protocol. Briefly, tissue was homogenized in 1 mL TRIzol™ with a bead homogenizer. After 5 min, the supernatant was mixed with 0.2 mL chloroform and centrifuged (30 min, 4°C, 12,000 × g). The aqueous phase was collected, mixed with 0.5 mL 2-propanol, incubated on ice for 10 min, and centrifuged (10 min, 4°C, 12,000 × g). RNA pellets were washed with 75% ethanol, centrifuged again, air-dried for 30 min, and dissolved in 60 µL RNase-free water at 55°C for 15 min. RNA concentration was measured using a NanoDrop™ 2000c (Thermo Fisher Scientific, Germany), and samples were stored at –80°C.

### Reverse Transcription and quantitative Real-Time PCR Analysis

One µg total RNA was used for cDNA synthesis using the SensiFAST™ cDNA Synthesis Kit (Meridian Bioscience Inc., USA) according to manufacturer’s instructions. The SensiFAST™ SYBR® No-ROX Kit (Meridian Bioscience Inc., USA) was used for quantitative real-time PCR analysis (T-Optical, Jena Analytics, Germany) with specific primers for the gene of interest listed in **Error! Reference source not found.**.

### Single Cell RNA Sequencing

Anti-GBM nephritis serum or PBS (control) was injected into mice, and kidneys were harvested at day 10 and 21 (n=3 mice per group/timepoint).

tdT-positive cells were isolated as previously described^26^. Briefly, cortical tissue was digested with collagenase type 1a (10 mg/ml, Life Technologies) and DNAse (10 mg/ml, Sigma Aldrich) for 40 min at 37°C. Afterwards, cells were filtered through a 40 µm nylon mesh (BD Bioscience), Fc-blocked (10 min, 4°C) and stained with hashtag antibodies (30 min, 4°C).

The tdTomato-positive cells were FACS-sorted and processed using Chromium Single Cell 3’ Library Kit (10x Genomics). Raw data were preprocessed with Cell Ranger. Quality control and downstream analysis were performed with Seurat (version 4.0) in R (version 4.2.2). Cells with <500 or>7000 genes or >15% mitochondrial content were excluded. In total, 18151 genes were detected. Data were normalized using *NormalizeData* function and highly variable features identified via *FindVariableFeatures (method = “vst”, nfeatures = 2000)*. Data were scaled using *ScaleData* with default parameters.

### Bioinformatic data analyses

#### Graph-based cluster analysis

Differentially expressed genes (DEGs) were identified using Seurat *FindAllMarkers* and ranked by log2 fold change. DEGs were visualized with *DoHeatmap*, ordered by fold change and displayed by genotype. Gene expression values were averaged per timepoint (*AverageExpression*). Compositional analyses were performed using *dplyr* (v1.5) and displayed as stacked bar plots.

#### Gene Ontology analysis

Upregulated genes in RenKO and WT samples were identified using relaxed thresholds to capture biologically relevant changes. Technical features (including cell hashing oligos, mitochondrial genes, ribosomal genes, and predicted genes) were excluded. Enrichment of biological process was performed using *clusterProfiler::enrichGO (organism = mouse)* with adjusted p-value <0.05 (Benjamini–Hochberg correction).

#### Pathway and Regulator Activity

Hallmark gene sets (MSigDB category H; *Mus musculus*) were retrieved using the *MSigDB* package and scored with *UCell*. Intracellular signaling pathway activity was inferred with *PROGENy* using normalized expression data from Seurat. Transcription factor activity was inferred using *DoRothEA* mouse regulons and the VIPER/decoupleR framework. Curated gene modules (interferon/STAT1, MAPK/AP-1, TGF-β effectors, fibroblast activation, and SMAD3 targets) were scored using UCell.

## Statistics

The data were visualized and analyzed using R (version 4.0.2)^27^ with RStudio (version 1.2.5033)^28^. A one-way ANOVA was performed to compare multiple groups, followed by Bonferroni multiple comparison test. P values <0.05 were considered statistically significant.

## Results

### Renin lineage cells are recruited to intraglomerular sites in crescentic glomerulonephritis

To investigate RLC involvement in cGN progression, we used the mRen-rtTAm2-LC1-tdT mouse line for inducible tdTomato-reporter tracing of RLCs. Following cGN induction, histological assessment was performed at days 10 and 21 (Fig. 1A). By day 10, crescents (≥2 cell layers) formed in 15% of glomeruli, accompanied by tuft hypercellularity, capillary collapse characterized by loss of capillary lumens, and extracellular matrix deposition (Fig. 1B). RLCs migrated from their juxtaglomerular position into glomeruli and along Bowman’s capsule, with 25% of glomeruli demonstrating overall RLC migration. Within the glomerular tuft, RLCs colocalized with α8 integrin-positive MCs and WT1-positive podocytes. Migration originated at the vascular pole, tracking crescent expansion toward the urinary pole. By day 21, disease progression was marked by persistent RLC migration in 20% of glomeruli, with RLCs localizing to regions of aberrant collagen-IV expression. These findings demonstrate sustained RLC recruitment throughout cGN progression, supporting a role of these cells in disease pathogenesis.

**Figure 1.**
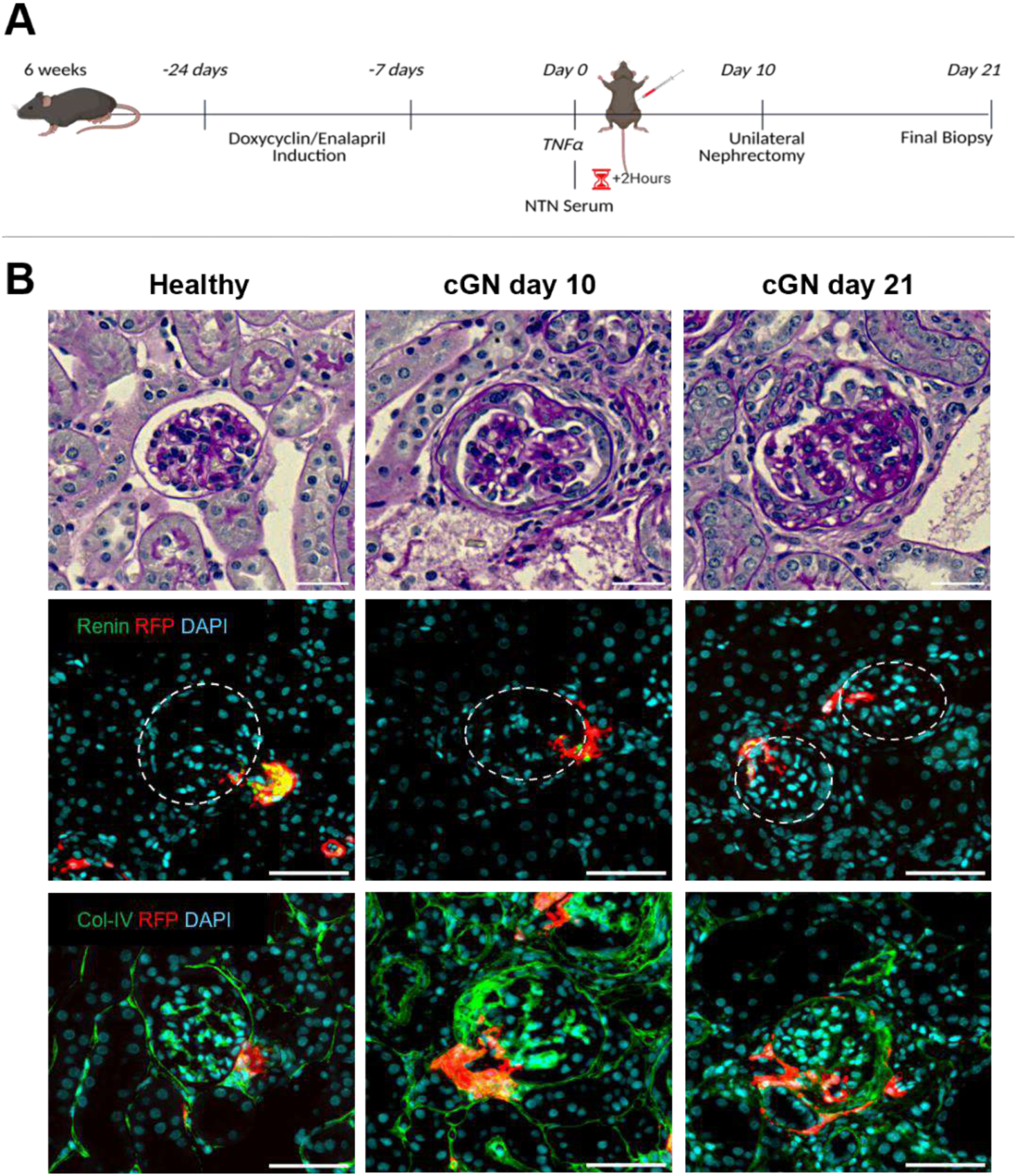
Renin Lineage cells localize to injured glomeruli in experimental crescentic glomerulonephritis (cGN). **A**) Experimental timeline of anti-GBM glomerulonephritis model induction. Kidney biopsies were collected at day 10 (unilateral nephrectomy) and day 21. **B**) First row: representative PAS-stained kidney sections showing crescent formation in wildtype mice at day 10 and day 21 compared to healthy control. Second row: Representative kidney sections stained for RFP (RLCs, red) and renin (green) and third row, stained for RFP (RLCs, red) and Col-lV (green) in wildtype mice at day 10 and day 21 compared to healthy control. Scale bar represents 50 µm. Dashed circles indicate glomerular boundaries. PAS, periodic acid-Schiff.

### Transgenic mice lead to expected phenotypes

To evaluate the role of renin production and RLCs themselves, two transgenic mouse lines were used. First, an RLC-specific renin knockout (RenKO) and an RLC ablation line via diphtheria toxin A (DTA) expression. Gene expression analysis showed significant renin reduction in both DTA and RenKO mice following genotype induction (Fig. 2A). Quantification of tdT⁺, renin⁺, and double-positive cells (Fig. 2B-D) further confirmed the expected phenotypes. RenKO mice retained tdT⁺ cells but showed ∼80% reduction in renin expression, whereas DTA mice exhibited 93% reduction in tdT⁺ cells and 70% reduction in renin⁺ cells. Both strains demonstrated significant reductions in tdT⁺/renin⁺ double-positive cells (96% in DTA, 94% in RenKO), confirming effective renin deletion or RLC ablation. The remaining renin⁺/tdT⁻ cells likely arise from incomplete recombination efficiency as previously reported, compensatory recruitment or neogenesis^12,29^.

**Figure 2.**
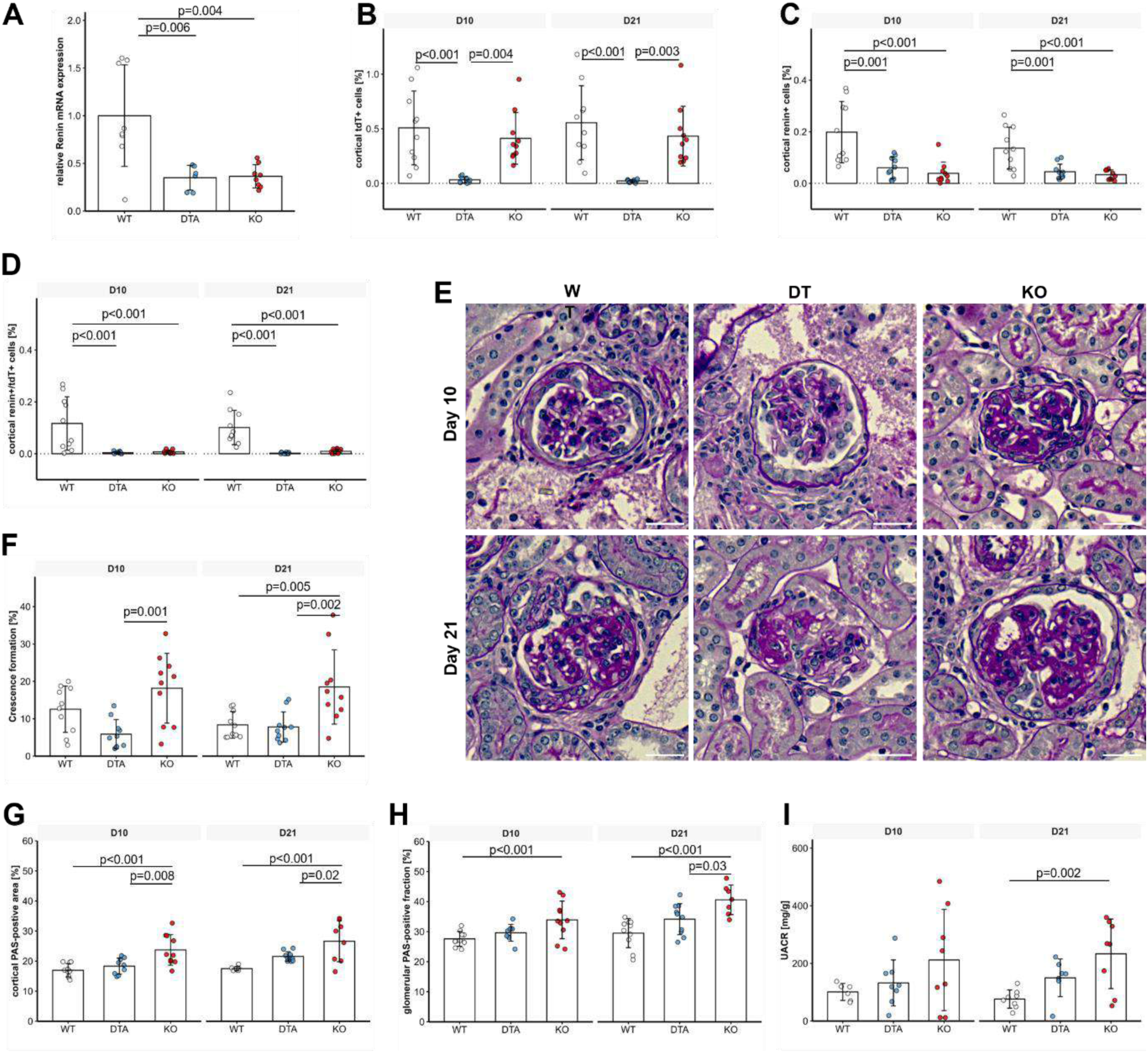
Ablation of RLCs or deletion of renin differentially influence the outcomes of crescentic glomerulonephritis. **A)** Relative renin mRNA expression in healthy WT, RenKO (KO), DTA mice to confirm genotype. **B)** Number of cortical tdT-positive cells, **C)** renin-positive cells, **D)** tdT⁺/renin⁺ double-positive cells in WT, RenKO, and DTA mice at day 10 and day 21. **E)** Representative PAS-stained kidney sections showing crescent formation at day 10 and day 21. **F)** Quantification of glomeruli presenting with crescent formation. **G)** Cortical and **H)** glomerular PAS-positive area [%]. **I)** Urinary albumin-to-creatinine (UACR) ratio [mg/g]. Number of animals 6-8 per group. Data shown as mean ± SD. Scale bar represents 50 μm. D10, day 10. D21, day 21. WT, wildtype. DTA, Diphtheria Toxin A RLC-ablation. KO, Renin Knockout.

Basal characterization of 3-month-old WT, RenKO, and DTA mice revealed no genotype-dependent differences in functional parameters (GFR, urine analysis, Supplemental Table 1) or histological markers (PAS, CD31, αSMA, Collagen IV, Supplemental Table 2).

### Renin KO but not DTA mice exhibited enhanced crescent formation

By day 10, DTA and RenKO groups displayed opposite trends in crescent formation relative to WT controls (Fig. 2F). DTA mice developed crescents in only 5.8% of glomeruli versus 12.5% in WT, whereas RenKO mice showed increased crescent formation (18.2% versus 12.5% in WT), accompanied by significantly elevated PAS-positive matrix accumulation (Fig. 2G). By day 21, RenKO mice showed significantly increased cortical and glomerular PAS-positive area and developed significantly more crescents relative to WT. While DTA mice did not show statistically significant differences in crescent formation or PAS positivity, they developed significant glomerular hypertrophy without increased glomerular cell number (Supplemental Fig. 1B–D).

To further characterize immune cell involvement in cGN, we quantified macrophage infiltration via CD68 staining (Fig. 3). On day 10, RenKO mice showed a trend toward increased macrophage infiltration compared to WT, though this did not reach statistical significance. Notably, RLC-depleted DTA mice exhibited significantly reduced macrophage infiltration compared to WT at both timepoints during cGN.

**Figure 3.**
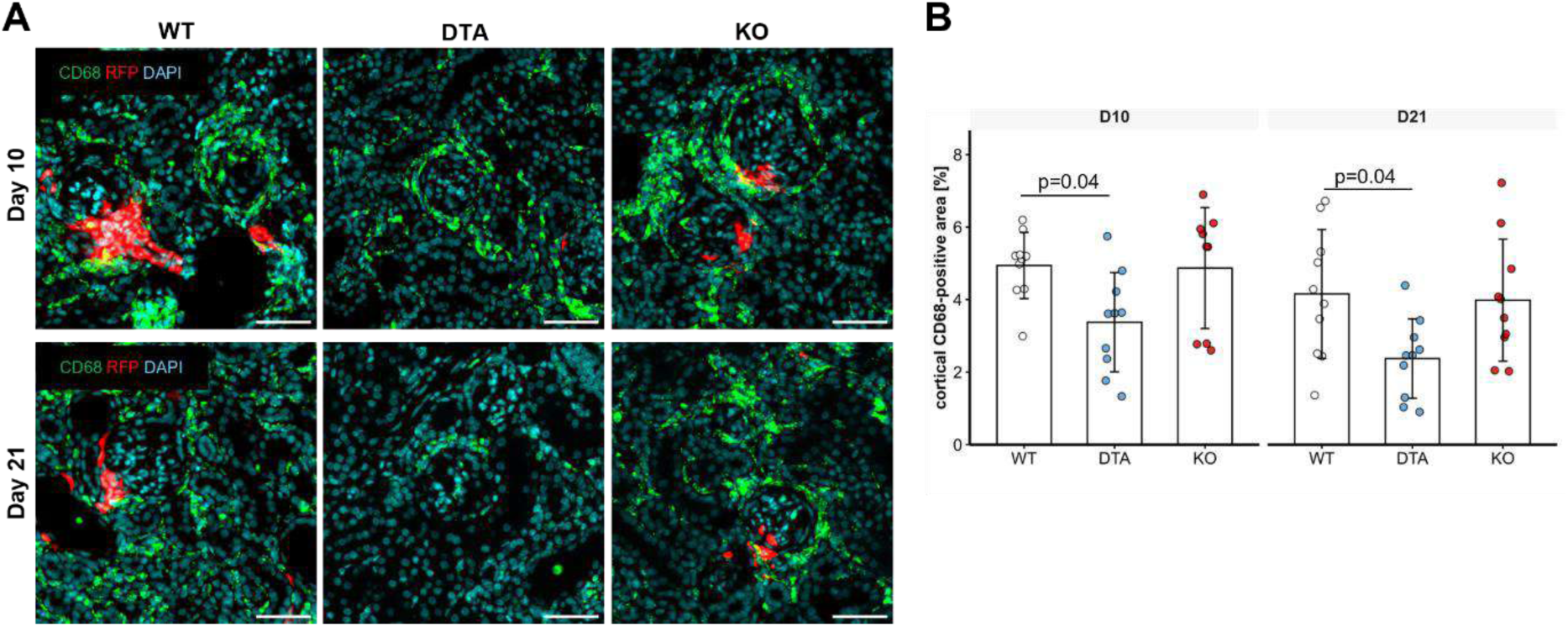
RLC ablation reduces CD68+ macrophage infiltration in crescentic glomerulonephritis. **A**) Representative images of CD68 (green) and tdTomato (red) staining at day 10 and day 21. Scale bar represents 50 µm **B**) Quantification of cortical CD68-positive area in WT, DTA, KO mice at day 10 and day 21. Dashed circles indicate glomerular boundaries. Data shown as mean ± SD.

To evaluate changes in glomerular cell composition and RLC-mediated regeneration, we assessed glomerular α8 integrin-positive area and the number of ERG- and WT1-positive cells (Fig. 4A-C). Mesangial cells (α8 integrin-positive area) and glomerular ERG-positive endothelial cells remained unchanged between groups at both timepoints. Podocyte number, quantified by WT1-positive staining, showed a trend toward decrease in RenKO mice compared to WT/DTA on day 10 and was significantly decreased on day 21 compared to both WT and DTA. Migration by tdT-positive cells was quantified by counting glomeruli with intraglomerular tdT-positive signal compared to the total number of glomeruli (Fig. 4D). The percentage of glomeruli with RLC migration was decreased in RenKO compared to WT on day 10 (9,2 % vs. 23,1%) and day 21 (10.0 % vs. 21.0%).

**Figure 4.**
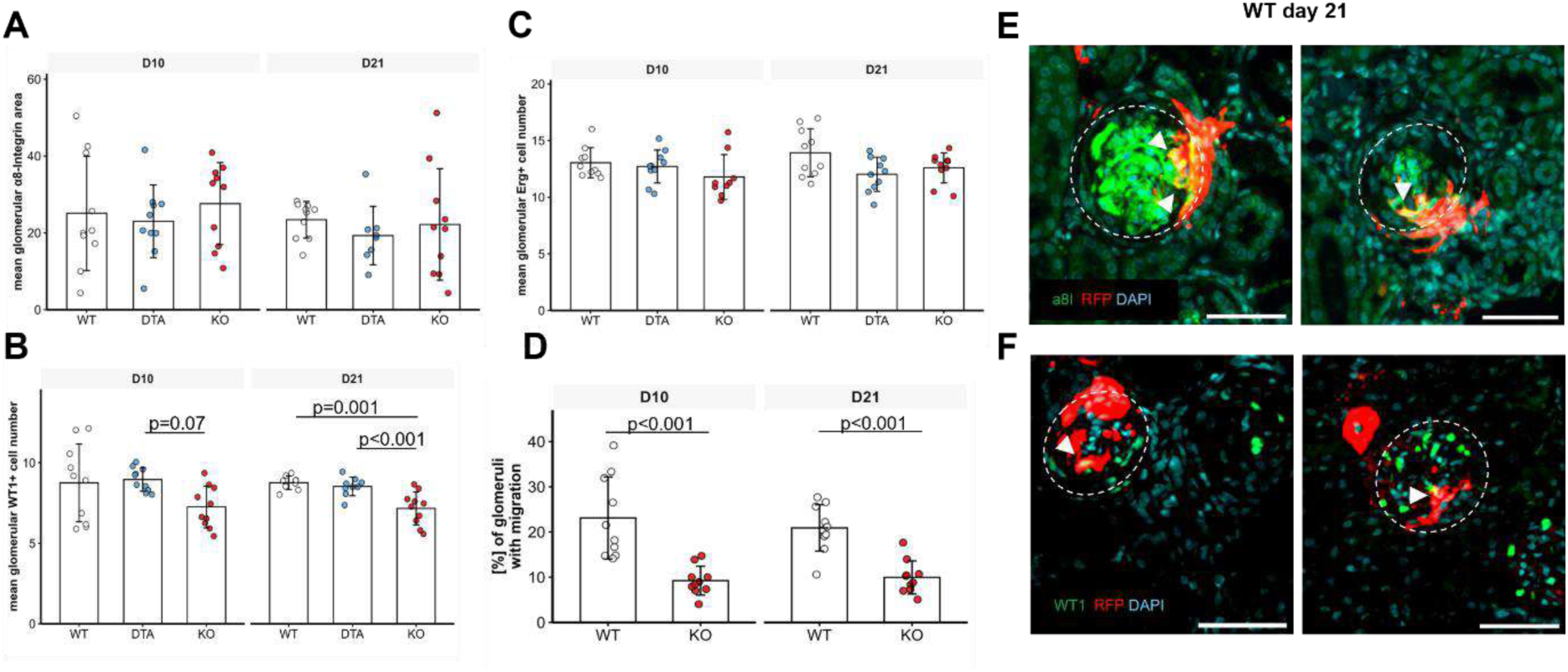
A) Quantification of mean a8-Integrin-positive area in glomeruli, **B)** mean number of glomerular WT1-positive cells and **C)** mean number of glomerular Erg-positive cells in WT, DTA and RenKO (KO) mice at day 10 and day 21. **D)** Quantification of glomeruli with intraglomerular tdT-positive signal at day 10 and day 21 in WT and RenKO mice. **E)** Representative immunofluorescence images showing overlapping (white arrow) α8-integrin (green), renin lineage cells (RFP, red), and DAPI (cyan) in WT glomeruli at day 21. Dashed circles indicate glomerular boundaries. **F)** Representative images showing WT1 (green), RLCs (RFP, red), and DAPI (cyan) overlapping in glomeruli at day 21. Scale bars = 50μm. Data shown as mean ± SD. n ≥ 9 animals per group (except α8I/DTA/day 21, n = 8).

We next examined whether renin expression affects RLC migration efficiency during cGN. While the number of tdTomato⁺ glomeruli in the JGA was similar between genotypes, the proportion that successfully migrated into injured glomeruli was significantly impaired in RenKO mice. At days 10 and 21, RLC migration occurred in 44-46% of tdT⁺ glomeruli in WT mice but only 18-23% in RenKO mice (Table 3). Analysis of migration patterns revealed additional differences. WT mice showed balanced distribution between intracapillary and extracapillary RLC migration, whereas RenKO mice exhibited predominant extracapillary migration (61% at day 21 versus 44% in WT) and reduced intracapillary migration (26% versus 41% in WT). Consistent with impaired intracapillary migration, RenKO mice showed significantly fewer RLCs colocalizing with MCs and podocyte markers (Table 3). These data suggest that renin expression is required for efficient RLC migration and intraglomerular cell replacement during glomerular injury.

**Table 1.**
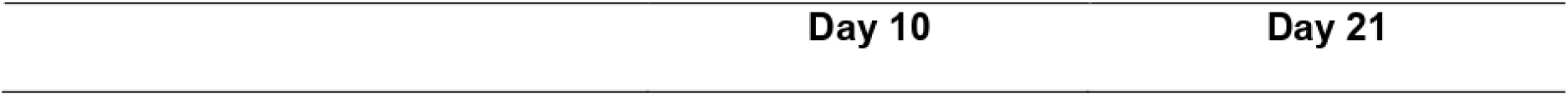
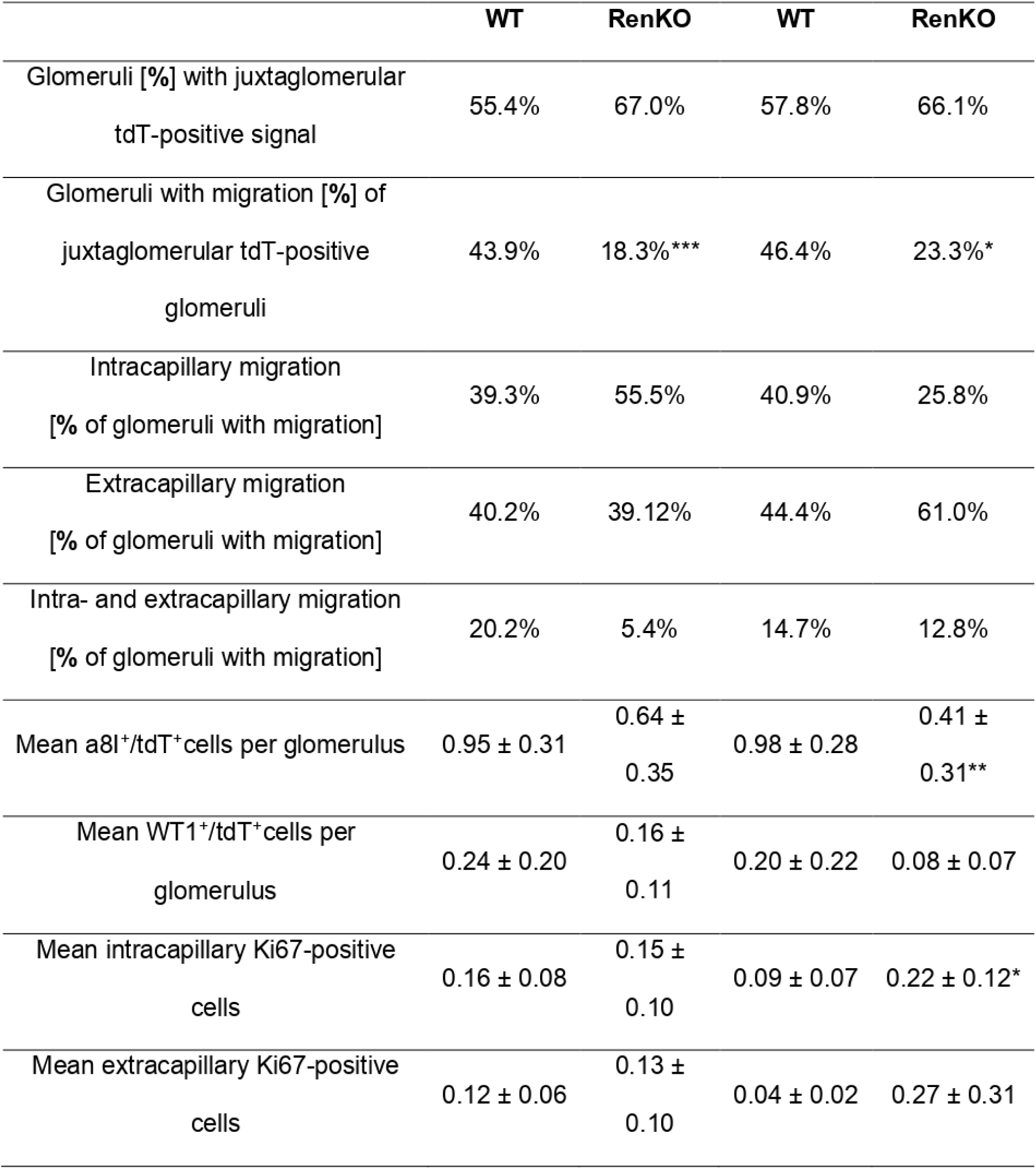
Quantification of RLC migration in wildtype (WT) and Renin-floxed (RenKO) mice. For statistical difference a t-test was performed (WT vs. RenKO at each timepoint). **\***p<0.05, **p<0.01, ***p<0.001.

|  | WT | RenKO | WT | RenKO |
| --- | --- | --- | --- | --- |
| Glomeruli [%] with juxtaglomerular tdT-positive signal | 55.4% | 67.0% | 57.8% | 66.1% |
| Glomeruli with migration [%] of juxtaglomerular tdT-positive glomeruli | 43.9% | 18.3%*** | 46.4% | 23.3%* |
| Intracapillary migration [% of glomeruli with migration] | 39.3% | 55.5% | 40.9% | 25.8% |
| Extracapillary migration [% of glomeruli with migration] | 40.2% | 39.12% | 44.4% | 61.0% |
| Intra- and extracapillary migration [% of glomeruli with migration] | 20.2% | 5.4% | 14.7% | 12.8% |
| Mean a8I<sup>+</sup>/tdT<sup>+</sup> cells per glomerulus | 0.95 ± 0.31 | 0.64 ± 0.35 | 0.98 ± 0.28 | 0.41 ± 0.31** |
| Mean WT1<sup>+</sup>/tdT<sup>+</sup> cells per glomerulus | 0.24 ± 0.20 | 0.16 ± 0.11 | 0.20 ± 0.22 | 0.08 ± 0.07 |
| Mean intracapillary Ki67-positive cells | 0.16 ± 0.08 | 0.15 ± 0.10 | 0.09 ± 0.07 | 0.22 ± 0.12* |
| Mean extracapillary Ki67-positive cells | 0.12 ± 0.06 | 0.13 ± 0.10 | 0.04 ± 0.02 | 0.27 ± 0.31 |

### Lack of renin leads to a different transcriptional profile in healthy mice

To characterize transcriptional changes in RLC lacking renin expression, we performed single-cell RNA sequencing on tdTomato-positive cells isolated from WT and RenKO mice under healthy conditions and following induction of cGN. Differential gene expression (DEG) analysis between WT and RenKO mice revealed marked differences in tdT-positive cells (Supplement Fig. 2A). Subsequent GO Biological Process enrichment analysis identified distinct molecular programs between genotypes in healthy mice.

In tdT-positive cells from healthy WT mice, which are derived solely from the JGA, the most significantly enriched categories were associated with RNA processing and mitochondrial function (Supplement Fig. 2B). Top enriched terms included RNA splicing, mRNA splicing via spliceosome, and regulation of mRNA metabolic processes, reflecting active transcriptional and post-transcriptional regulation. In parallel, we observed strong enrichment of pathways linked to mitochondrial organization and energy metabolism, including *oxidative phosphorylation*, *cellular respiration*, and *aerobic respiration*.

In contrast, the tdT-positive cells of RenKO mice displayed a different enrichment profile (Supplement Fig. 2C). Top categories were related to cytoskeletal reorganization and cell signaling, including *actin filament organization*, *regulation of actin cytoskeleton organization*, and *small GTPase-mediated signal transduction*. Moreover, tdT-positive cells with renin deletion exhibited upregulation of pathways associated with protein stability and catabolism, along with pronounced enrichment of immune-related processes, such as *antigen processing and presentation via MHC class II*. This immunomodulatory profile suggests that renin-deficient RLC may acquire antigen-presenting capabilities, potentially contributing to local immune surveillance or inflammatory responses in the kidney.

### Lack of renin impairs the ability of renin cells for cell migration

Next, tdT-positive cells were isolated from WT and RenKO mice after cGN induction and analyzed at two time points. The analysis revealed pronounced differences between genotypes (Fig. 5). Volcano plots show the differentially expressed genes (DEGs) at day 10 and day 21 (Fig. 5A, B), with markedly increased differential expression at day 21 compared to day 10. A heatmap of the top 30 DEGs at D21 is shown in (Fig. 5D).

**Figure 5.**
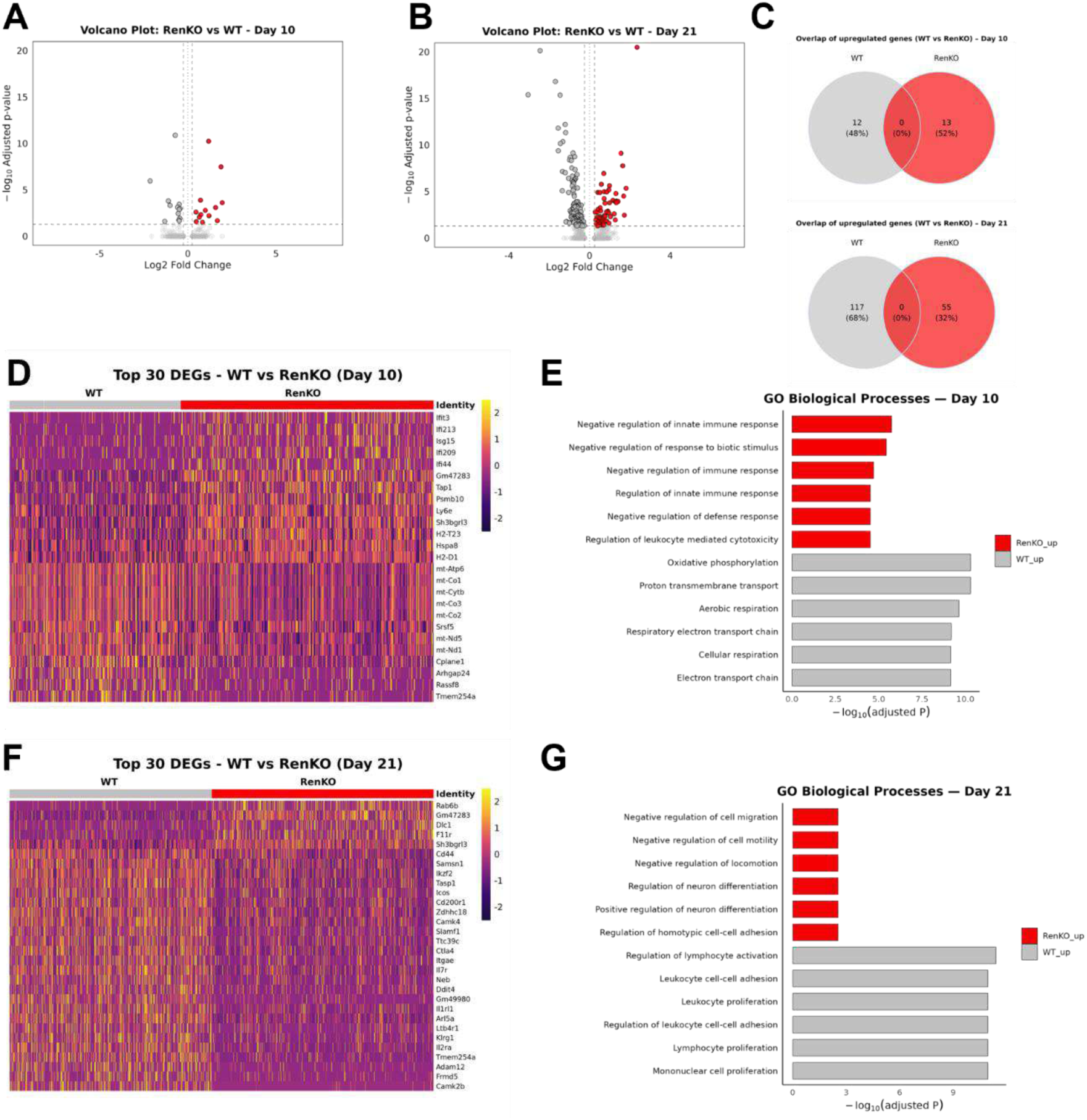
Renin deletion drives transcriptional reprogramming of Renin lineage cells in crescentic glomerulonephritis. **A)** Volcano plots of differentially expressed genes (DEGs) in tdT-positive RLCs comparing RenKO versus WT mice at day 10 and **B)** day 21. Red dots indicate significantly upregulated genes; gray dots represent non-significant changes. **C)** Venn diagrams showing overlap of upregulated genes between WT and RenKO at day 10 (top) and day 21 (bottom). **D)** Heatmap of the top 30 DEGs between WT and RenKO at day 10 and **E)** Gene Ontology (GO) biological process enrichment analysis of DEGs at day 10. **F)** Heatmap of the top 30 DEGs between WT and RenKO at day 21 and **G)** GO analysis of DEGs at day 21. Bar length represents −log10 (adjusted p-value); red bars indicate pathways enriched in RenKO; gray bars indicate pathways enriched in WT.

Gene Ontology (GO) enrichment analysis revealed that renin deficiency alters the metabolic and immune profiles of RLCs. At day 10, while WT tdT-positive cells exhibited enrichment of categories associated with oxidative phosphorylation, transmembrane transport, and aerobic and cellular respiration, indicating elevated metabolic activity consistent with increased energy demand during the early immune response (Fig. 5E), renin-deficient RLC failed to upregulate these metabolic programs. Instead, RenKO mice showed significant enrichment of GO categories related to negative regulation of innate immune responses and responses to biotic stimuli, suggesting that loss of renin alters immunoregulatory programs in RLCs (Fig. 5F).

At D21, while WT tdT-positive cells displayed significant enrichment of GO categories associated with leukocyte cell-cell adhesion, leukocyte proliferation, and T-cell activation, reflecting ongoing immune cell activation and recruitment (Fig. 5G), renin deficiency impaired these processes. RenKO tdT-positive cells instead showed enrichment of terms related to negative regulation of cellular movement, including cell migration, motility, and locomotion. This transcriptional state is consistent with the impaired migratory behavior of RenKO RLC observed histologically and may limit their capacity to participate in reparative responses during glomerular injury.

Additionally, pathway interference revealed distinct genotype-specific differences in tdT-positive cell signaling programs during cGN. JAK/STAT signaling activity was significantly elevated in RenKO at both day 10 and day 21 (Fig. 6A), consistent with enhanced interferon-driven responses (Fig. 6B), suggesting a dysregulated inflammatory state characterized by persistent type I interferon signaling. DoRothEA transcription factor (TF) activity analysis further supported these findings, showing that STAT1 was selectively upregulated in the tdT-positive cells of RenKO mice at day 10 (Fig. 6D). MAPK pathway activity showed a modest but significant increase in RenKO mice, especially at day 21 (Fig. 6A), suggesting heightened stress-related transcriptional responses during the late inflammatory phase.

**Figure 6.**
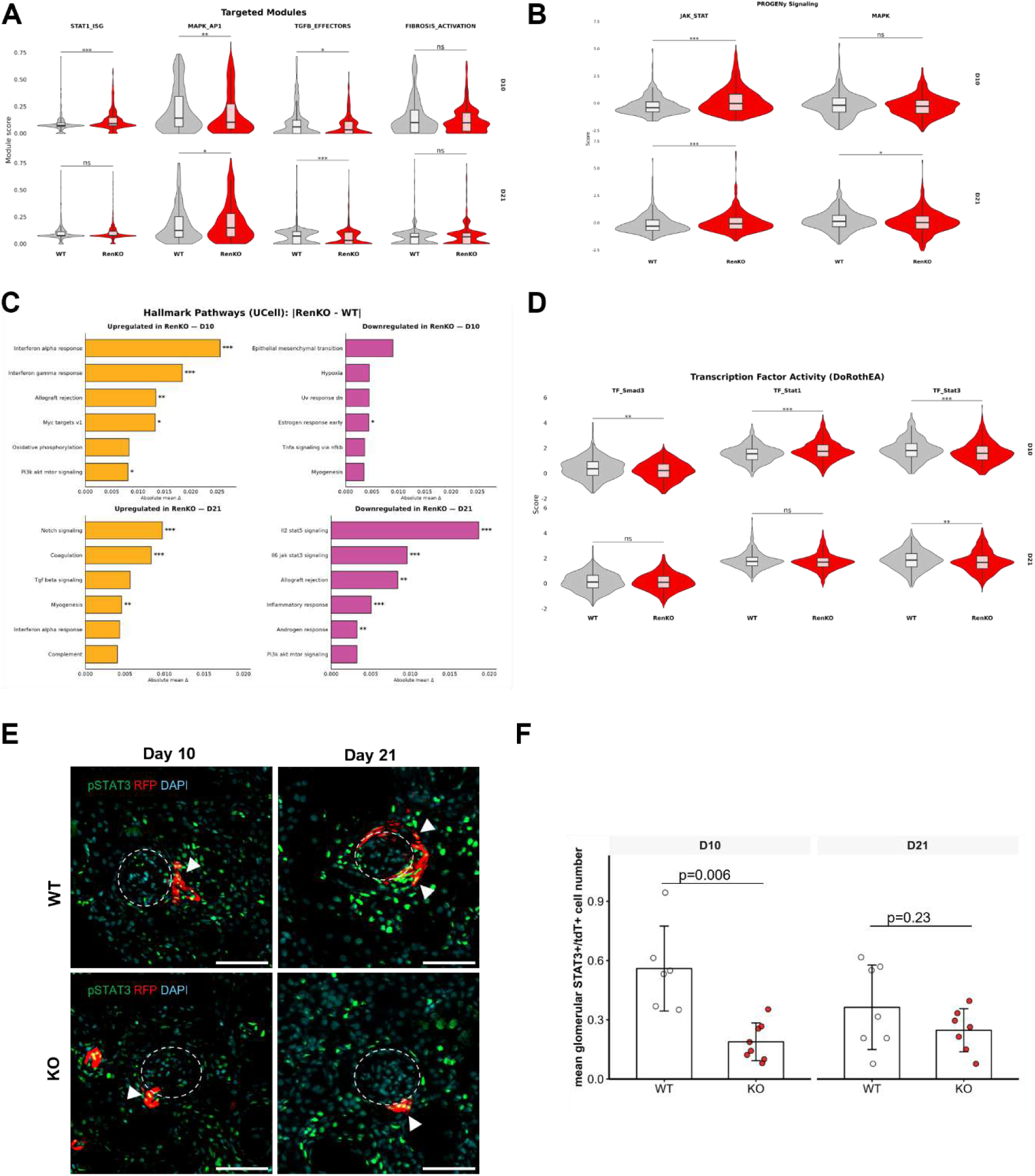
Renin deletion associates with Renin lineage cells shift from immune-fibrotic (STAT3/SMAD3) to inflammatory (STAT1) signaling. **A)** Violin plots of targeted module scores for STAT1, MAPK, TGF-β effectors, and fibroblast activation in tdT-positive cells of WT and RenKO at day 10 and day 21. **B)** PROGENy pathway activity scores comparing JAK-STAT and MAPK signaling between WT and RenKO across timepoints. **C)** Hallmark pathway activity (UCell) showing differential enrichment in tdT-positive cells of RenKO versus WT at day 10 (top) and day 21 (bottom). Orange bars indicate pathways upregulated in RenKO; purple bars indicate pathways downregulated in RenKO. **D)** DoRothEA-inferred transcription factor activity for SMAD3, STAT1, and STAT3 in WT and RenKO tdT-positive cells at day 10 and day 21. **E)** Representative immunofluorescence images showing phospho-STAT3 (pSTAT3, green), tdTomato-labeled RLCs (red), and DAPI (blue) in kidney sections from WT and RenKO (KO) mice at day 10 and 21. Dashed white circles outline glomeruli. White arrows indicate pSTAT3+RFP+ double-positive cells. Scale bar = 50 μm. **F)** Quantification of STAT3 activation showing the ratio of pSTAT3+RFP+ double-positive cells to total tdTomato+ cells per glomerulus at day 10 and 21. Data shown as mean ± SD.

Conversely, TGFβ pathway activation, quantified both by PROGENy scores and targeted TGFB_EFFECTORS module scoring, was consistently lower in tdT-positive cells of RenKO, indicating impaired canonical SMAD3-mediated responses in RenKO, but preserved signaling in WT RLC (Fig. 6A, C). This attenuated TGFβ signaling in RenKO may disrupt critical cellular responses required for proper glomerular repair and homeostasis. In addition to lower SMAD3 activity in RenKO tdT-positive cells at day 10, STAT3 activity remained decreased in RenKO cells at both day 10 and day 21, highlighting a disrupted IL6–STAT3/SMAD3 axis in RenKO versus a sustained axis in WT RLCs (Fig. 6D, E).

### Renin deficiency impairs STAT3 phosphorylation in juxtaglomerular RLCs

To validate the transcriptional finding of impaired RLC STAT3 signaling at the protein level, we performed immunofluorescence staining for phosphorylated STAT3 (pSTAT3) at days 10 and 21 (Fig. 6E). pSTAT3-positive RLCs were predominantly localized to the JGA in both genotypes and time points. At day 10, RenKO mice demonstrated a significant reduction in the proportion of pSTAT3+tdTomato+ double-positive cells compared to WT (Fig. 6F) consistent with impaired STAT3 activation in renin-deficient RLCs. The absence of a significant difference at day 21 suggests that STAT3 activation in RLC is temporally restricted to the early injury phase.

## Discussion

In this study, we investigated the differential effects of RLC ablation versus RLC-specific renin knockout in murine cGN. Unexpectedly, renin deficiency in juxtaglomerular RLCs more prominently altered disease progression than RLC ablation. While RLC ablation reduced macrophage infiltration and increased glomerular hypertrophy with subtle effects on crescent formation, RenKO mice developed aggravated pathology by day 21, with increased albuminuria, crescent formation, PAS positivity, and podocyte loss. Suggesting a causative relationship, RenKO RLCs migrated into injured glomeruli significantly less than WT RLCs at both timepoints. Single-cell analysis revealed that transcriptional plasticity toward migratory/regenerative but also and inflammatory programs of RLC is renin-dependent.

Using lineage tracing in mRen-rtTAm2-LC1-tdT mice, we showed that juxtaglomerular-derived RLCs participate in both intraglomerular inflammation and regeneration, extending prior work in regenerative injury models^10–12,29,30^. During cGN, migrated RLCs localized to α8 integrin⁺ MCs, WT1⁺ podocytes, and along Bowman’s capsule, with sporadic incorporation into crescentic layers. This data confirms that juxtaglomerular RLCs can serve as progenitors for MCs^12^, PECs and podocytes^11^ supporting repair but also show for the first time that they are able to contribute to crescent formation and cGN progression either directly or via paracrine effects.

Despite substantial RLC depletion, DTA mice displayed only modest disease changes and reduced macrophage infiltration, contrasting with the marked effects of renin deficiency. Several factors may contribute to this finding. First, RLC ablation was incomplete (∼10% of cells remaining). Second, even in WT mice, only one-quarter of glomeruli showed RLC migration histologically, corresponding to roughly half of glomeruli analyzed using 3D methodology^12^. Third, our data provide evidence for compensatory recruitment of alternative renin-producing cells. Despite ∼90% depletion of tdT⁺ RLCs in our DTA mice, cortical renin⁺ cells were reduced by only 60-70%, and renin mRNA by 60% This discordance indicates non-lineage-traced cell populations contribute to compensatory renin synthesis. This compensation is specific to the DTA model, as RenKO mice retain RLCs comparable in number to WT despite genetic deletion of renin. Several renin-competent cell types could serve as compensatory sources. Vascular smooth muscle cells (VSMCs) retain epigenetic memory of the renin phenotype and can be recruited for renin synthesis^31^, while extraglomerular MCs or pericytes can produce renin under stress, a reversible shift consistent with broader plasticity in the renin-producing system^32^. Moreover, neogenesis persists in the adult kidney, whereby cells from non-renin-expressing populations can undergo *de novo* differentiation into renin-producing cells when homeostasis is threatened^29,31,33^. Thus, the renin-producing system exhibits plasticity through VSMCs recruitment, MC conversion, and neogenesis, collectively explaining the phenotypic similarity between DTA and WT despite near-complete RLC ablation.

Our findings suggest that adult RLCs not only contribute to regeneration but also participate in inflammatory responses driving disease progression. RLCs localized to or within crescents, and scRNA-seq revealed that WT RLCs display an immune-enriched transcriptomic landscape with differential expression of cytokine receptors (Il1rl1, Il7r, Il2ra), co-stimulatory molecules (Cd44, Ctla4, Icos, Slamf1), and enrichment of immune-related GO terms (WT, day 21). We identified Rab6b among the differentially expressed genes, previously shown to be induced in glomerular cells via TNFR1 signaling and implicated in trafficking and secretion of inflammatory mediators^34^. However, comparable disease progression in DTA and WT emphasizes that crescent formation is orchestrated by diverse cellular pathways beyond RLC-mediated effects on inflammation or PEC activation^35–38^. These findings are consistent with observations in mice, where Gsα/cAMP-defective RLCs induced microvascular kidney injury through phenotypic alteration. These findings parallel observations in healthy mice where Gsα/cAMP-defective RLCs induced microvascular injury through phenotypic alteration. This suggests that changes in RLC phenotype, rather than loss, may be pivotal for modulating disease under severe pathological stress^21^.

In contrast to mild effects in RLC-ablated mice, renin deletion in adult RLCs markedly worsened cGN. Compared to WT, RenKO animals developed higher proteinuria, increased crescent percentage, greater PAS positivity, and podocyte loss. While prior work established a pathogenic role of the Ang II–AT1R axis, showing that AT1R inhibition reduces crescent formation, it may appear paradoxical that renin loss worsens crescentic GN^39,40^. However, this paradox may be resolved by considering that renin deletion reduces not only Ang II–AT1R activity but also substrates for protective pathways such as AT2R and ACE2–Ang-(1–7)–Mas known to preserve podocytes and limit fibrosis^41–43^. Moreover, intrarenal AngII can be generated via renin-independent proteases (chymase, cathepsin G), maintaining AT1R signaling^44–46^. Thus, renin loss during cGN may create imbalance favoring pathogenic over protective signaling. Our study data are also consistent with reports demonstrating that during nephrogenesis the Ang2-AT1R axis is indispensable relating to the frequent finding for many systems that important embryonic mechanisms might be switched on in adult life in regenerative/protective programs after organ injury^47^.

Beyond RAS imbalance, renin expression may preserve RLC phenotypic identity and migratory/regenerative ability. Previous work demonstrated that renin deletion drives RLCs towards a matrix-secretory phenotype, promoting vascular disease^21,48^. Consistent with this, we observed markedly reduced RLC migration in RenKO mice, implicating renin production as an important factor of RLCs migration to glomeruli upon injury.

In addition to our histological findings, scRNA-seq analysis also revealed that renin deletion altered RLC phenotype in cGN. At day 10, RenKO RLCs failed to upregulate metabolic programs supporting migration and repair. While renin-competent WT RLCs increased expression of genes driving ATP generation through oxidative phosphorylation (mt-Atp6, mt-Co1, mt-Co2, mt-Co3, mt-Cytb), processes linked to migration, stemness, and proliferation^49^, RenKO RLCs mounted an elevated interferon-stimulated gene response (Ifit3, Isg15). This maladaptive interferon-driven program is recognized as proinflammatory and compromises repair capacity in cGN^38^. The pathogenic role of ISG responses has been documented in murine as well as human cGN, where interferon activation in infiltrating and resident podocytes and PEC exacerbates injury^50,51^.

By day 21, transcriptional divergence became more pronounced. RenKO RLCs exhibited enrichment of genes involved in negative regulation of cell migration and locomotion, indicating loss of migratory capacity and adoption of a structurally anchored phenotype. In contrast, renin-competent WT RLCs maintained expression of pathways related to T-cell activation and adhesion, including immune checkpoint molecules Ctla4 and Cd200r1^52,53^. While classically associated with T-cell regulation, their expression in progenitor populations may represent a mechanism through which RLCs influence local immune responses during severe cGN inflammation.

Pathway analysis revealed that renin deletion shifts RLC signaling. RenKO RLCs exhibited a Stat1/interferon-driven inflammatory signaling associated with maladaptive responses and progression to chronic kidney disease^54–57^. In contrast, WT RLCs activated Stat3/Smad3-dependent pathways during the critical early injury phase. Immunofluorescence validation confirmed significantly reduced STAT3 phosphorylation in RenKO RLC at day 10, providing protein-level evidence for the transcriptional impairment of the IL6-STAT3 axis. While STAT3 signaling can be pathogenic when chronically sustained, it has been shown to promote epithelial survival, controlled matrix repair, and coordination of inflammatory responses in acute injury settings^58–61^. The early STAT3 activation in WT RLC may therefore represent an adaptive response that enables migration and balanced immune modulation. The failure of renin-deficient RLC to mount this early STAT3 response, coupled with elevated JAK/STAT1 activity, creates a pathological imbalance in which reparative signaling programs are not properly coordinated. This is further supported by the loss of migratory capacity and adoption of an interferon-driven, structurally anchored phenotype by day 21 in RenKO mice. Thus, renin expression appears to preserve an adaptive reparative phenotype early in injury, whereas renin deficiency constrains RLCs into a maladaptive inflammatory state characterized by impaired STAT3-dependent repair mechanisms.

Several limitations call for consideration. Neither RLC ablation nor renin deletion achieved complete 100% efficiency. While renin deletion markedly influenced disease outcomes, compensatory mechanisms in DTA mice may have mitigated more pronounced effects. Second, although our model is well-established for murine cGN, it does not fully recapitulate human crescentic disease heterogeneity, and direct clinical translation requires caution. Third, while we establish that renin deletion worsens cGN and impairs RLC migration, the precise mechanisms remain to be fully elucidated. Nevertheless, the genetic specificity of our model with renin deletion exclusively in RLCs, provides a foundation for future mechanistic studies, and the integration of lineage tracing, functional outcomes, and single-cell transcriptomics provides multi-level evidence for renin-dependent RLC plasticity in cGN.

In summary, adult RLCs contribute to murine cGN as both precursors replacing injured glomerular cells and modulators of local inflammation. RLC ablation produced mild anti-inflammatory effects without altering outcome, suggesting compensatory mechanisms. In contrast, renin production emerged as critical for RLC identity and functional versatility. In WT mice, RLCs maintained migratory capacity, metabolic function, immune checkpoint molecule expression supporting repair and immune modulation, whereas renin deletion constrained RLCs into an interferon-driven, non-migratory state with impaired reparative capacity and exacerbated injury.

In conclusion these data indicate that RLC plasticity, and their balance between reparative and inflammatory phenotypes, is tightly linked to renin expression. Targeted modulation of renin-dependent RLC plasticity may therefore represent a promising, cell-specific strategy for treating glomerulonephritis.

## Disclosures

The authors declare that there are no conflicts of interest regarding the publication of this paper.

## Funding

This work was supported by Deutsche Forschungsgemeinschaft (DFG) Projekt TO 679/5-1 (470138795) (to V.T.) and HU 600/8-1.

## Acknowledgements

We are very grateful to Ulf Panzer and Anett Peters for providing the NTS serum and for their valuable advice. We also highly acknowledge the excellent technical assistance of Anika Wirth, Andrea Angermann, Katja Maiwald, and Maria Schuster.

## Author Contributions

S.A. performed experiments, analyzed and interpreted data, performed single-cell RNA-sequencing analysis, wrote the original draft, edited the manuscript; S.H. performed experiments, acquired and interpreted data; A.S. performed experiments, acquired and analyzed data; A.A. and I.S. performed experiments; A.D. and R.G. performed single-cell RNA-sequencing library preparation and 10x Genomics-based transcriptomic experiments; J.S. and F.G. provided conceptualization and supervision; V.T. acquired funding and reviewed the manuscript; S.B. provided manuscript supervision; H.W. provided supervision, analyzed data, wrote and edited the manuscript; C.H. acquired funding, provided conceptualization and supervision, reviewed and edited the manuscript.

## Data Sharing Statement

The data supporting the findings of this study are available from the corresponding author upon reasonable request.

## Notes

### Competing Interest Statement

The authors have declared no competing interest.

