## Supplemental Files for "Renin is critical for Renin Lineage Cell Plasticity and Migration in experimental crescentic Glomerulonephritis"

**Supplement – Table of Content**

**Additional Materials & Methods**

**Supplement Table 1:** Characterization of kidney function in wildtype (WT), Renin-floxed (KO) and Renin cell-ablated (DTA) mice.

**Supplement Table 2:** Characterization of kidney morphology from wildtype (WT), Renin-floxed (KO) and Renin cell-ablated (DTA) mice.

**Supplement Figure 1:** Additional characterization of kidney morphology and function in wildtype (WT), Renin-floxed (KO) and Renin cell-ablated (DTA) mice.

**Supplement Figure 2:** Single cell gene expression analysis of healthy wildtype (WT) and Renin-floxed (KO) mice.

**Additional Materials & Methods**

**Animals – Generation of KO mice**

The RenKO strain was generated in cooperation with Prof. Frank Schweda (University of Regensburg)^19^. Therefore, mouse embryonic stem (ES) cells with floxed renin allele were obtained from EUCOMM (Project 40265, Ren1tm1a(EUCOMM) Wtsi). Ren1c floxed mice were generated by the injection of ES cells in blastocysts of C57/BL6 mice at the Transgenic Core Facility, Max Planck Institute of Molecular Cell Biology and Genetics (MPI-CBG), Dresden. The resulting chimeras were propagated to obtain mice homozygous for the renin floxed allele. Results were confirmed by PCR. We then crossed these mice to the mRen-rtTAm2-LC1-tdT strain.

**Immunofluorescence stainings**

Supplement Table 1: List of primary and secondary antibodies.

| Anti- | Contributor | Reference number |
| --- | --- | --- |
| RFP | Biorbyt | orb182397 |
| Renin | abcam | ab212197 |
| alpha-smooth muscle actin | abcam | ab7817 |
| Integrin alpha-8 | Santa Cruz Biotechnology | sc-365798 |
| WT-1 | abcam | ab89901 |
| ERG | abcam | ab92513 |
| Goat-IgG, AF647 | ThermoFisher | A-21447 |
| Mouse-IgG, AF488 | ThermoFisher | A-21202 |
| Rabbit-IgG, AF555 | Abcam | ab150074 |

Quantitative Real-Time PCR Analysis

Supplement Table 2: Primer pairs used for quantitative PCR analysis.

| **Target** | **Forward** | **Reverse** |
| --- | --- | --- |
| **Renin** | 5’-GCTACATGGAGAACGGGTCC-3’ | 5’-CCATGCCTAGAACACCGTCA-3’ |
| **Hprt** | 5’-CGAGGAGTCCTGTTGATGTTGC-3’ | 5’-CTGGCCTATAGGCTCATAGTGC-3’ |

**Supplement Table 1: Characterization of kidney function in wildtype (WT), Renin-floxed (KO) and Renin cell-ablated (DTA) mice.** After induction of recombination with doxycycline-containing drinking water for 3 weeks, mice underwent Uninephrectomy. One week later kidney function was analyzed by measuring glomerular filtration rate (GFR) and analysis of 24h urine collected in metabolic cages.

|  | **WT** | **KO** | **DTA** |
| --- | --- | --- | --- |
| **Body weight**  [g] | 28.7 ± 3.5 | 29.4 ± 3.9 | 25.8+3.6 |
| **Water intake**  [ml/24h] | 1.1 ± 0.7 | 1.1 ± 0.2 | 1.1+0.7 |
| **Urine volume**  [ml/24h] | 1.7 ± 0.8 | 1.7 ± 0.8 | 1.2+0.5 |
| **Water balance**  [ml/24h] | -0.6 ± 0.8 | -0.6 ± 0.6 | -0.1+0.9 |
| **Sodium excretion**  [µmol/24h] | 184.4 ± 64.3 | 182.6 ± 54 | 134.5+58.5 |
| **Potassium excretion** [µmol/24h] | 390.5 ± 144.4 | 300.8 ± 86.6 | 377.5+126.2 |
| **Urine osmolality**  [mmol/kg water] | 1944.4 ± 642.8 | 1625.6 ± 674.2 | 2488.0+936.8 |
| **Osmolyte excretion**  [µmol/24h] | 2912.8 ± 1011.2 | 2384.7 ± 594.0 | 2611.1+794.6 |
| **Creatinine excretion** [µmol/24h] | 5.1 ± 2.2 | 5.1 ± 1.9 | 4.3+1.4 |
| **Albumin excretion**  [µmol/24h] | 35.3 ± 17.5 | 23.9 ± 11.7 | 26.4+15.0 |
| **Albumin-creatinine-ratio**  [A.U.] | 6.8 ± 1.9 | 1.0 ± 0.4 | 6.1+3.0 |
| **GFR**  [µl/min/100g b.w.] | 541.3 ± 93.6 | 602.4 ± 106.6 | 692.6+139.9 |

**Supplement Table 2: Characterization of kidney morphology from wildtype (WT), Renin-floxed (KO) and Renin cell-ablated (DTA) mice.** After induction of recombination with doxycycline-containing drinking water for 3 weeks, mice underwent uninephrectomy. Histological samples were stained and semi-automatically analyzed.

|  | **WT** | **KO** | **DTA** |
| --- | --- | --- | --- |
| **CD31**  [% of positive kidney area] | 4.7 ± 0.6 | 4.8 ± 0.6 | 4.7 ± 0.7 |
| **Collagen IV**  [% of positive kidney area] | 1.5 ± 0.9 | 0.8 ± 0.5 | 0.9 ± 0.8 |
| **α-SMA**  [% of positive kidney area] | 0.6 ± 0.2 | 0.6 ± 0.2 | 0.7 ± 0.2 |
| **PAS positivity**  [% of positive kidney area] | 15.3 ± 3.2 | 16.7 ± 1.5 | 16.9 ± 4.0 |
| **Glomerular PAS positivity**  [% of positive glomerular area] | 25.4 ± 5.6 | 30.1 ± 3.1 | 27.9 ± 6.5 |
| **Mean glomerular area**  [µm²] | 1869.2 ± 205.5 | 1991.1 ± 134.0 | 1764.9 ± 130.1 |
| **Glomerular nuclear number**  [mean number of nuclei per glomerulus] | 26.6 ± 2.4 | 27.8 ± 1.5 | 25.8 ± 1.1 |
| **Glomerular nuclear density**  [mean number of nuclei per µm² of glomerular area] | 0.014 ± 0.001 | 0.014 ± 0.001 | 0.015 ± 0.001 |

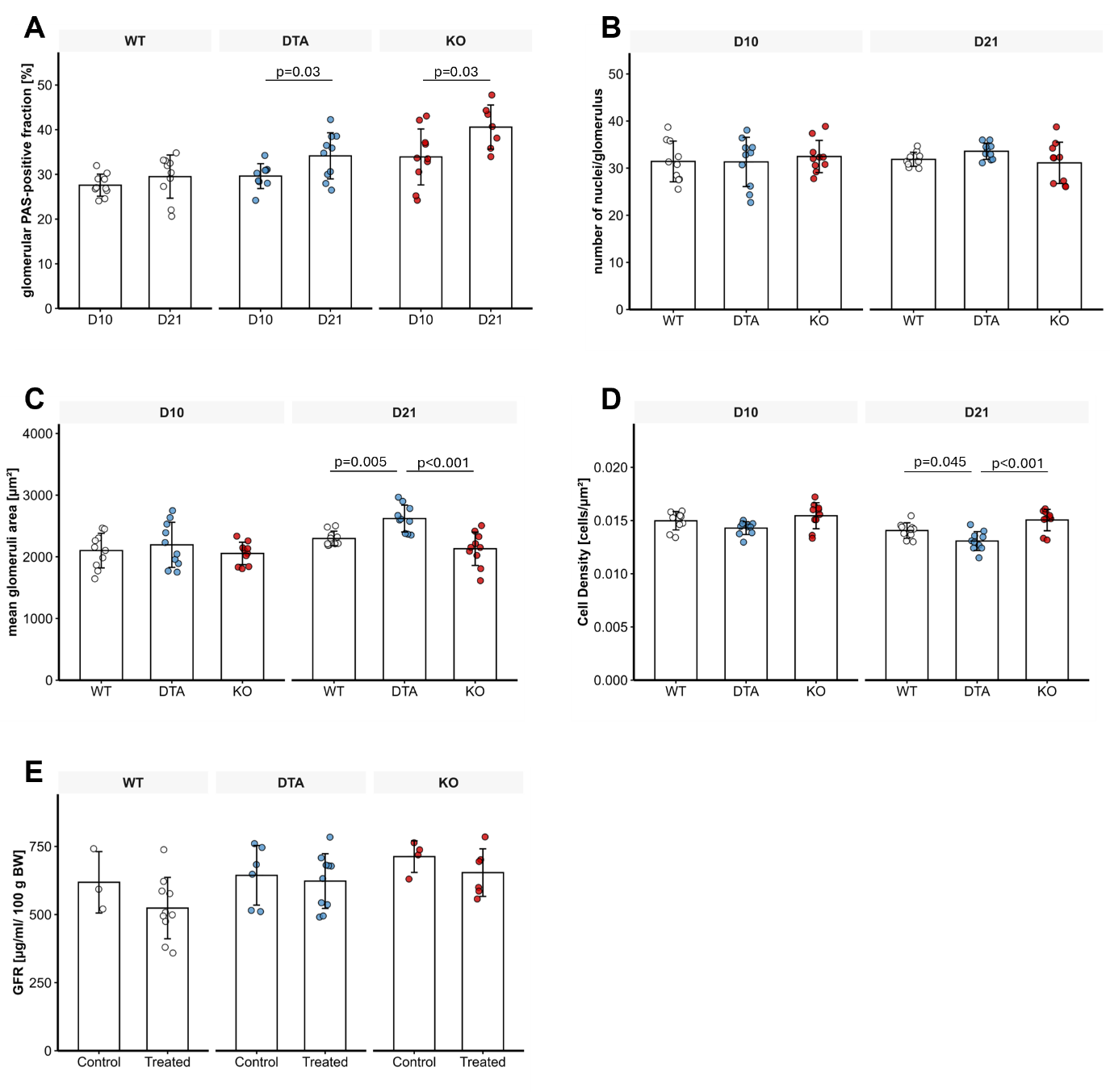

**Supplement Figure 1:** **Additional characterization of kidney morphology and function A)** Time-dependet comparison of glomerular PAS-positive area. **B)** Average number of nuclei per glomerulus. **C)** Mean glomerular area and **D)** Mean glomerular cell density in WT, DTA and KO mice at D10 and D21. Number of animals 10 in each group. **E)** Glomerular filtration rate (GFR) of healthy control and anti-GBM treated WT, DTA and KO mice at D21. Number of animals 3-6 for healthy control mice and 6-10 for anti-GBM treated mice.

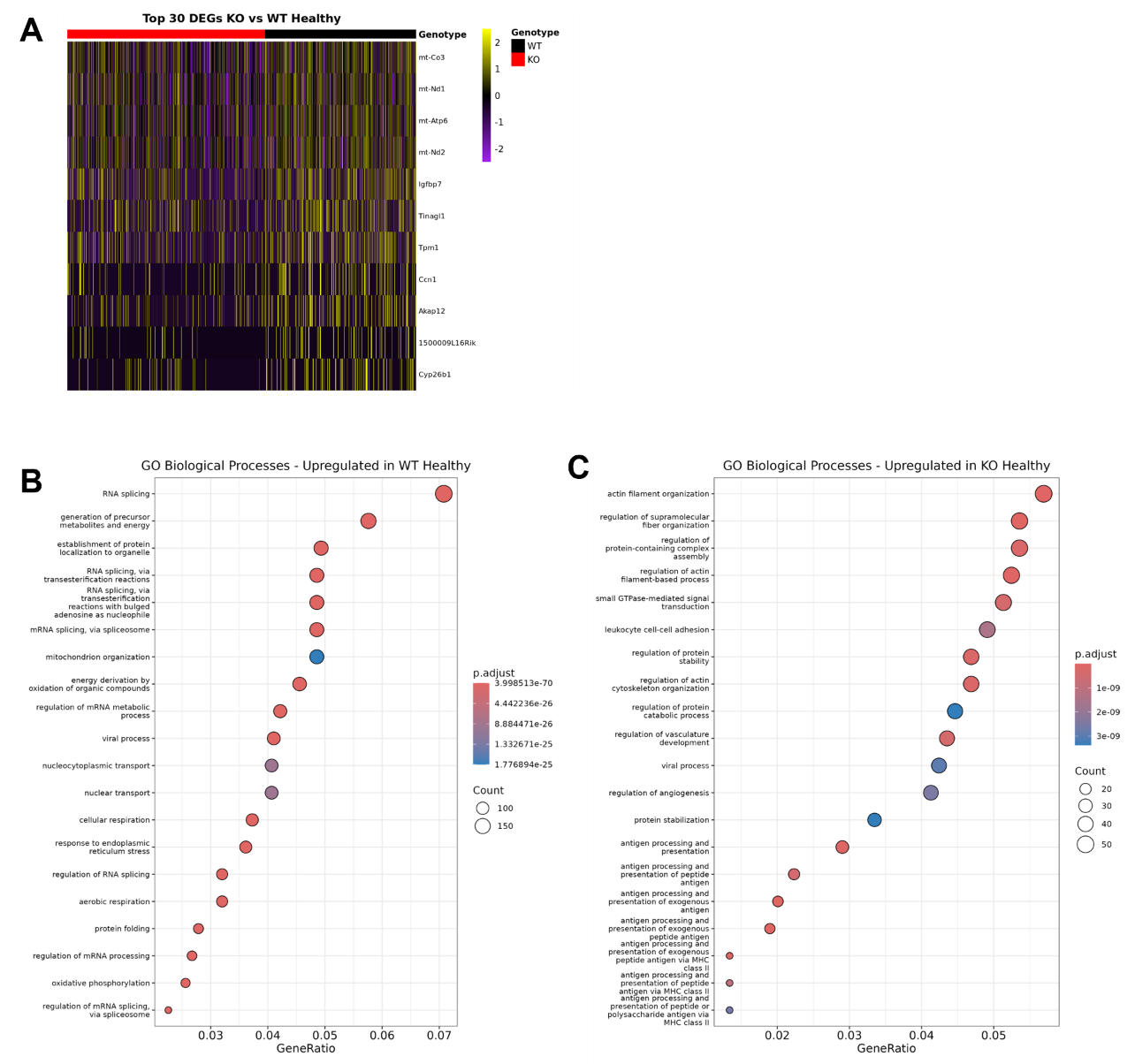
**Supplement Figure 2 A)** Heatmap analysis with DEGs between healthy WT and KO mice. **B)** The most enriched 20 categories identified by GO analysis of DEGs of WT (left) and KO (right).
